# Oxytocinergic modulation of stress-associated amygdala-hippocampus pathways in humans is mediated by serotonergic mechanisms

**DOI:** 10.1101/2021.11.06.467580

**Authors:** Chunmei Lan, Congcong Liu, Keshuang Li, Zhiying Zhao, Jiaxin Yang, Yina Ma, Dirk Scheele, Yao Shuxia, Keith M. Kendrick, Benjamin Becker

**Affiliations:** The Clinical Hospital of the Chengdu Brain Science Institute, School of Life Science and Technology, University of Electronic Science and Technology of China, 611731 Chengdu, China; Department of Psychology, Xinxiang Medical University, Henan 453003, China; School of Psychology and Cognitive Science, East China Normal University, 200062 Shanghai, China; Department of Radiology and Biomedical Imaging, Yale University School of Medicine, New Haven, Connecticut; State Key Laboratory of Cognitive Neuroscience and Learning, IDG/McGovern Institute of Brain Research, Beijing Normal University, 100875 Beijing, China; Division of Medical Psychology, Department of Psychiatry and Psychotherapy, University Hospital Bonn, 53105 Bonn, Germany; Department of Psychiatry, School of Medicine & Health Sciences, University of Oldenburg, 26129 Oldenburg, Germany

**Keywords:** oxytocin, serotonin, amygdala, stress, anxiety

## Abstract

**Background:** The hypothalamic neuropeptide oxytocin (OXT) may exert anxiolytic and stress-reducing actions via modulatory effects on amygdala circuits. Animal models and initial findings in humans suggest that some of these effects are mediated by interactions with other neurotransmitter systems, in particular the serotonin (5-HT) system. Against this background, the present pharmacological resting state fMRI study aimed at determining whether effects of OXT on stress-associated amygdala intrinsic networks are mediated by 5-HT.

**Methods:** We employed a randomized placebo-controlled double-blind parallel-group pharmacological fMRI resting state experiment during which n = 112 healthy male participants underwent a transient decrease in 5-HT signaling via acute tryptophan depletion (ATD) or the corresponding placebo-control protocols (ATDc) before the administration of intranasal OXT or placebo intranasal spray, respectively.

**Results:** OXT and 5-HT modulation exerted interactive effects on the coupling of the left amygdala with the ipsilateral hippocampus and adjacent midbrain. OXT increased intrinsic coupling in this pathway, while this effect of OXT was significantly attenuated during transiently decreased central serotonergic signaling induced via ATD. In the absence of OXT or 5-HT modulation this pathway showed a trend for an association with self-reported stress perception in everyday life. No interactive effects were observed for the right amygdala.

**Conclusions:** Together, the findings provide first evidence that effects of OXT on stress-associated amygdala-hippocampal-midbrain pathways are critically mediated by the 5-HT system in men.

## Introduction

The hypothalamic neuropeptide oxytocin (OXT) regulates socio-emotional behavior across species, with convergent evidence from experimental studies in humans suggesting that the intranasal administration of OXT can e.g. exert pro-social, anxiolytic and stress-reducing effects (Heinrichs et al., 2009; Meyer-Lindenberg et al., 2011; Yao et al., 2018; Xin et al., 2020; Grinevich and Neumann, 2021; Quintana et al., 2021). Accumulating evidence from sophisticated animal models suggests that some of the complex regulatory effects of OXT are critically mediated by interactions with other neurotransmitter systems, such that animal models demonstrated that dopamine partly mediated OXT’s effects on pair bonding (Young and Wang, 2004) while the serotonin (5-HT) system critically mediated OXT’s regulatory effect in the domains of social reward and anxiety (Yoshida et al., 2009; Dölen et al., 2013; Lefevre et al., 2017). In terms of the anxiolytic properties, a previous rodent study demonstrated a dense expression of OXT receptors in the raphe nucleus, the primary source of central 5-HT and reported that the administration of exogenous OXT facilitated 5-HT release in this region and subsequently reduced anxiety-like behavior (Yoshida et al., 2009). Based on these findings, we recently examined the interactive effects between OXT and 5-HT in humans on threat-related amygdala activity using a randomized parallel-group placebo-controlled pharmacological functional magnetic resonance imaging (fMRI) experiment which combined the intranasal administration of OXT with an acute tryptophan depletion (ATD) procedure (Liu et al., 2021a). The administration of an ATD procedure represents a robust means to induce a transient decrease in central serotonergic signaling (Veen et al., 2007; Crockett et al., 2008; Evers et al., 2010), and thus pre-treatment with ATD may attenuate some of the potential 5-HT-mediated effects of OXT. We observed that OXT switched amygdala sensitization to social threat signals to desensitization and this potential anxiolytic action of OXT was attenuated following ATD-induced decreased central serotonergic signaling (Liu et al., 2021a).

The amygdala plays a crucial role in fear- and anxiety-related processes (e.g. (Phelps and LeDoux, 2005; Mihov et al., 2013); however, see also (LeDoux and Pine, 2016; Gothard, 2020; Zhou et al., 2021)) and both, OXT and 5-HT effects on the amygdala have been repeatedly documented (Cools et al., 2005; Fisher and Hariri, 2013; Kanat et al., 2015; Raab et al., 2016; Xin et al., 2020; Kreuder et al., 2020). OXT and 5-HT-sensitive receptors are widely distributed across limbic-striatal-frontal regions (Pasqualetti et al., 1998; Gimpl and Fahrenholz, 2001; Varnäs et al., 2004; Quintana et al., 2019) and thus may not only influence regional activity but also the functional cross-talk between nodes in these networks (Bethlehem et al., 2013). To examine modulatory effects of OXT on the network level while controlling for its valence- and context-specific effects (e.g. (Shamay-Tsoory and Abu-Akel, 2016; Yao et al., 2018; Chen et al., 2020)), an increasing number of studies employed pharmacological resting-state fMRI (pharmaco-rsfMRI) (see e.g. (Brodmann et al., 2017; Wu et al., 2020; Jiang et al., 2021; Xin et al., 2021)).

Previous studies employing placebo-control pharmaco-rsfMRI strategy to determine effects of the intranasal OXT on the intrinsic amygdala networks reported an OXT-induced modulation of the intrinsic coupling of the amygdala with prefrontal regions, particularly medial prefrontal (mPFC) and orbitofrontal regions (Fan et al., 2014; Dodhia et al., 2014; Eckstein et al., 2017; Cheng et al., 2018; Jiang et al., 2021), posterior default mode network (DMN) regions (Kumar et al., 2014) as well as with the hippocampal formation (Fan et al., 2015; Alaerts et al., 2019) with some studies reporting associations between intranasal OXT effects and stress- and emotion processing-associated indices including subclinical depressive symptoms or stress exposure (Fan et al., 2014, 2015; Eckstein et al., 2017). Although fewer studies employed a pharmaco-rsfMRI or ATD-rsfMRI approach to examine modulatory effects of 5-HT on the intrinsic amygdala networks some evidence indicates a serotonergic modulation of pathways partly overlapping with those observed following oxytocinergic modulation, including amygdala-posterior DMN (Dutta et al., 2019; Zhang et al., 2019), amygdala-prefrontal (Eisner et al., 2017) and amygdala-hippocampal connectivity (Carhart-Harris et al., 2015).

Further support for a potential network-level modulatory role of the 5-HT-OXT interactions comes from molecular imaging studies which demonstrated that the administration of exogenous OXT increased 5-HT concentrations in key limbic regions including the amygdala and hippocampus in non-human primates (Lefevre et al., 2017) and that intranasal OXT influenced serotonergic signaling in a broad network encompassing limbic, insular and prefrontal regions in humans (Mottolese et al., 2014; Lefevre et al., 2018). Importantly the study confirmed the central role of the amygdala in regulating 5-HT signaling via OXT such that OXT-induced 5-HT1A receptor binding changes in the amygdala which correlated with changes in the hippocampus, insula and prefrontal cortex (Mottolese et al., 2014). The identified amygdala networks partly resemble pathways involved in stress reactivity and emotion regulation (Etkin et al., 2015). In particular pathways such as the amygdala-hippocampal (Admon et al., 2009; Vaisvaser et al., 2013) and amygdala-prefrontal circuits are sensitive to both, long-term as well as acute stress-exposure (Herringa et al., 2013; Fan et al., 2015, 2014; Park et al., 2018).

Summarizing, accumulating evidence from animal and human models suggest an interaction of OXT and 5-HT in regulating anxiety- and stress-related behavior, however, while both systems have been associated with modulating the amygdala-centered networks, interactive effects on the intrinsic amygdala networks in humans have not been systematically examined. To this end we combined the administration of OXT or placebo (PLC) intranasal spray with an ATD (Acute tryptophan depletion) - induced transient decrease in central 5-HT signaling or a matched acute tryptophan depletion control (ATDc) protocol in a randomized controlled pharmaco-rsfMRI parallel group design in n = 121 healthy male subjects (details see also (Liu et al., 2021a)). Based on previous studies, we hypothesized that OXT’s effects on the amygdala intrinsic networks are (partly) mediated by interaction with the 5-HT system and that a transient reduction in 5-HT signaling following ATD would attenuate OXT’s effect, and (2) given the role of OXT and 5-HT in stress processing and a high stress-sensitivity of the amygdala intrinsic networks (Sripada et al., 2012; Vaisvaser et al., 2013; Fan et al., 2015; Zhang et al., 2016; Feng et al., 2018) the identified pathways would be associated with levels of currently experienced stress exposure.

## Material and methods

### Participants

A total of 121 non-smoking, right-handed, young healthy male participants were enrolled (detailed study enrollment criteria see also (Liu et al., 2021a)). Given that previous studies reported sex-differences with respect to both, the effects of OXT on amygdala functional connectivity (Ma et al., 2018) and central 5-HT synthesis rates (Nishizawa et al., 1997) and to further control for potential confounding effects of OXT administration with hormonal changes across the menstrual cycle the present study focused on male individuals (similar approach see also Eckstein et al., 2017; Zhao et al., 2019a; Xin et al., 2020). Participants were instructed to abstain from alcohol and caffeine for 24 hours and from food and drinks (except water) for 12 hours prior to the experiment. To examine interaction effects between the OXT and 5-HT systems, a randomized double-blind placebo-controlled between-group pharmacological resting state fMRI (pharmaco-rsfMRI) design was employed during which the subjects received ATD or an acute tryptophan depletion control (ATDc) drink which balanced for tryptophan before they were administered either OXT or a corresponding placebo (PLC) intranasal spray and subsequently underwent resting state fMRI. During initial quality assessments data from 9 subjects were excluded from the following analyses due to mania or depression history (n = 2), technical failure of the MRI system (n = 1), poor spatial registration quality (n = 1) and excessive head motion > 3mm (n = 5) (details see CONSORT flowchart Supplementary **Figure S1**) leading to a total number of n = 112 subjects for the final analysis (ATD-OXT, n = 29; ATD-PLC, n = 29; ATDc-OXT, n = 26; ATDc-PLC, n = 28, detailed group characteristics see **Table 1**).

**Table 1.** Group characteristics of subject age and questionnaire scores.

| Measurements | ATD-OXT<br>N = 29 | ATD-PLC<br>N = 29 | ATDc-OXT<br>N = 26 | ATDc-PLC<br>N = 28 | $F_{(3,108)}$ | $p$ |
| --- | --- | --- | --- | --- | --- | --- |
| Age(years) | 22.0 (2.4) | 21.9 (2.6) | 22.2 (2.3) | 22.1 (2.2) | 0.09 | 0.97 |
| BDI | 6.0 (6.3) | 7.0 (5.7) | 5.2 (6.6) | 5.1 (5.0) | 0.66 | 0.58 |
| STAI-SAI | 37.7 (8.9) | 34.8 (6.3) | 34.9 (8.2) | 35.9 (6.7) | 0.87 | 0.46 |
| STAI-TAI | 41.1 (8.3) | 40.5 (5.9) | 38.4 (7.5) | 41.2 (6.8) | 0.88 | 0.45 |
| PSS | 14.2 (5.5) | 14.9 (4.7) | 13.8 (4.9) | 15.5 (4.7) | 0.62 | 0.61 |
| PANAS-P(T1) | 24.9 (6.9) | 23.9 (8.3) | 24.3 (6.9) | 25.1 (8.2) | 0.15 | 0.93 |
| PANAS-N(T1) | 15.7 (9.0) | 13.1 (5.7) | 11.9 (5.2) | 13.3 (6.0) | 1.56 | 0.20 |

**Abbreviations:** OXT, Oxytocin; PLC, Placebo; ATD, Acute Tryptophan Depletion; ATDc, Acute Tryptophan Depletion control mixture; BDI, Beck's Depression Inventory; STAI-TAI, State-Trait Anxiety Inventory-Trait Anxiety Inventory; STAI-SAI, State-Trait Anxiety Inventory-State Anxiety Inventory; PSS, Perceived Stress Scale; PANAS-P, Positive and Negative Affect Schedule-Positive affect; PANAS-N, Positive and Negative Affect Schedule-Negative affect; T1, pre-oral administration.

All participants provided written informed consent after being fully informed about the experimental procedures. The study had full ethical approval from the Ethics Committee at the University of Electronic Science and Technology of China and adhered to the latest revision of the declaration of Helsinki. The study was preregistered on ClinicalTrials.gov (NCT03426176, https://clinicaltrials.gov/show/ NCT03426176).

### Procedure

Experimental and treatment procedures were identical to our previous study which reports the results of a task-based fMRI paradigm in the same sample to assess interactive effects of oxytocin and serotonin on amygdala threat reactivity and sensitization (Liu et al., 2021a). Briefly, a randomized, double-blind, placebo-controlled between-subject pharmaco-rsfMRI design was employed during which four treatment groups received combinations of amino acid mixture drinks (ATD vs. ATDc) to induce a transient decrease in central serotonergic signaling and intranasal spray (OXT vs. PLC) to modulate central oxytocin signaling. To adhere to the pharmacodynamic profile of treatments participants arrived between 7:30 to 10:00 AM and underwent fMRI acquisition between 13:30 to 16:00 PM. Upon arrival, participants received a standardized protein-poor diet for breakfast. Following the assessment of pre-treatment control variables, participants underwent a previously validated tryptophan depletion protocol with ATD (detailed components of mixtures see Supplementary **Table S1**) which has been demonstrated to lead to a robust transient reduction in central 5HT signaling (Veen et al., 2007; Crockett et al., 2008; Passamonti et al., 2012) or ATDc. The ingestion of the amino acid mixtures was followed by a resting period of 5 hours to achieve a robust reduction in tryptophan levels. During the resting period participants were asked to relax and magazines were provided. Subsequently, control variables were re-assessed and participants administered either OXT (24IU, ingredients: oxytocin, glycerine, sodium chloride and purified water) or PLC (identical ingredients except for oxytocin) intranasal spray. Both intranasal sprays were provided by Sichuan Meike Pharmaceutical Co. Ltd in China. In line with the pharmacokinetic profile of intranasal OXT (Spengler et al., 2017b), rsfMRI data was collected 45 min after OXT/PLC administration. Control variables were re-assessed before and after fMRI acquisition (schematic outline of the experimental protocols see **Figure 1**). The rsfMRI acquisition was followed by a task-based fMRI paradigm assessing amygdala threat reactivity and sensitization (Liu et al., 2021a).

**Figure 1.**
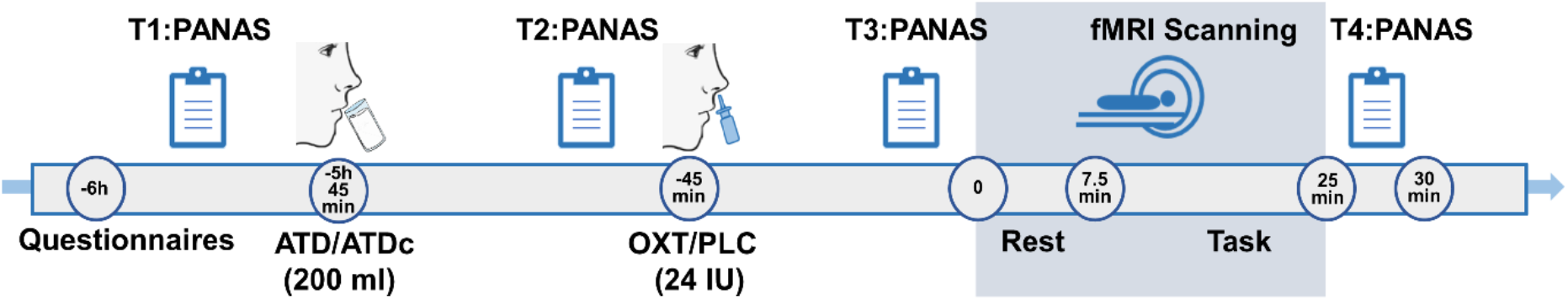
Experimental protocols and procedures of the pharmacological resting-state fMRI experiment.

### Behavioral assessments

Anxiety and depression have been associated with amygdala functional networks (Spengler et al., 2017a; Zhao et al., 2019b; Xu et al., 2021; Liu et al., 2021b) and therefore corresponding indices were assessed to control for pre-treatment differences between the groups. To this end the State-Trait Anxiety Inventory (STAI) (Spielberger et al., 1970) and Beck Depression Inventory (BDI-II) (Beck et al., 1996) were applied. The Positive and Negative Affect Schedule (PANAS) (Watson et al., 1988) was repeatedly administered before administration of the amino acid drink (T1) and the intranasal spray (T2) as well as immediately before MRI acquisition (T3) and at the end of the experiment (T4) to control for unspecific effects of treatment on mood (procedure see **Figure 1**). No between-group differences in the potential confounders were observed (details see **Table 1**).

Both OXT and 5-HT have been strongly associated with stress processing and stress reactivity (Olff et al., 2013; Mahar et al., 2014) as well as intrinsic amygdala functional networks (e.g. Seeley et al., 2018; Zhang et al., 2019; Jiang et al., 2021). However, the intrinsic amygdala networks have been associated with numerous functional domains, including not only acute stress (Archer et al., 2018), but also e.g. trait anger (Fulwiler et al., 2012), discrimination (Clark et al., 2018), social functioning (Johns et al., 2019), or early life stress exposure (Luo et al., 2022). We therefore included a measure of perceived stress during the previous month using the Perceived Stress Scale (PSS) (Cohen et al., 1983) to explore whether the identified pathways are associated with current stress. To control for treatment effects on the assessment and the neural pathway the questionnaire was applied before treatment and the analysis focused on the subjects without active treatment.

### MRI data acquisition

MRI data was obtained on a 3-T GE MR750 Discovery MRI system (General Electric Medical System, Milwaukee, WI, USA). High-resolution brain structural data were acquired with a T1-weighted sequence using the following parameters: repetition time, 6.0 ms; echo time, 1 ms; flip angle, 12°; field of view, 256×256 mm; resolution, 256×256; slice thickness, 1 mm; number of slices, 156. Resting-state functional MRI data were acquired using an echo planar imaging sequence with the following acquisition parameters: repetition time, 2000 ms; echo time, 30 ms; field of view, 220 × 220 mm; flip angle, 90°; resolution, 64×64; slice thickness, 3.2 mm; number of slices, 43. A total number of 225 whole-brain volumes were collected (about 7.5 min). During scanning, two head cushions were used to prevent excessive head motion while ensuring comfort. Participants were instructed to relax and think of nothing in particular while fixating a fixation cross presented centrally via a rear mirror.

### Functional MRI data preprocessing

The first 10 volumes of the resting-state time-series were discarded for each subject and subsequent fMRI data preprocessing was carried out using FSL FEAT 6.0 (https://www.fmrib.ox.ac.uk/fsl). The following preprocessing steps were applied: brain extraction with BET (Brain Extraction Tool, v2.1), motion correction, slice-timing correction, spatial smoothing with a Gaussian kernel of 5 mm full width at half maximum, intensity normalization and 0.01-0.1 Hz band-pass filtering after ICA-AROMA was employed (Pruim et al., 2015). Mean signals from white matter and cerebrospinal fluid (CSF) were removed by means of linear regression. Registration of functional data to high resolution structural images was carried out using FLIRT with boundary-based registration (Jenkinson and Smith, 2001; Jenkinson et al., 2002; Greve and Fischl, 2009). Registration from high resolution structural to standard space was then further improved using FNIRT nonlinear registration with 12 degrees of freedom. To control for the critical impact of head motion on functional connectivity analyses (Van Dijk et al., 2012), ICA-AROMA was applied to remove motion-related artifacts before filtering, subjects with head motion > 3 mm or high mean frame-wise displacement (mean FD, > 0.3 mm) were excluded, and for the group-level analysis (Power et al., 2012) mean FD was included as a covariate. Kruskal-Wallis one way-ANOVA indicated that there was no significant difference between the four treatment groups with respect to mean FD (FD_ATD-OXT_=0.1, *SD*=0.05, FD_ATD-PLC_ = *SD*= 0.03, FD_ATDc-OXT_=0.08, *SD* =0.02, FD_ATDc-PLC_=0.07, *SD* = 0.03, *F* _(3,108)_ =6.65, *p* > 0.05).

### Functional connectivity analyses

In line with our research question, the main and interaction effects of treatments (ATD and OXT vs. the respective control treatment conditions) were examined on the amygdala intrinsic networks by means of a seed-to-whole brain resting state connectivity analysis. Anatomical masks of the left amygdala and right amygdala from the Automated Anatomic Labeling (AAL) served as a priori defined seed regions. Individual functional connectivity (FC) maps were initially created for each participant using DPASFA 4.4 (advanced edition of DPABI, http://rfmri.org/dpabi) and each seed region by calculating Pearson correlations between the mean time-course extracted from the amygdala masks and all other voxels in the brain and subsequently transformed to z-maps using Fisher r-to-z transformation.

Effects of treatment were examined separately for the left and right amygdala intrinsic connectivity networks by means of a 2 × 2 ANCOVA in SPM12 (Friston et al., 1994) with amino acid mixture (ATD/ATDc) and intranasal spray (OXT/PLC) as between-subjects factor and mean FD and age as covariates and the grey matter mask template from DPABI as explicit mask. Our main hypothesis in terms of interactive effects between the OXT and 5-HT system was evaluated by means of applying a whole brain analysis with a stringent initial cluster-forming threshold which combine voxel wise *p* < 0.001 with cluster wise *p* < 0.05 family-wise error (FWE) corrected (Woo et al., 2014; Eklund et al., 2016; Daniel Kessler, 2017). The probabilistic maps from the Anatomy toolbox (version 2.2) (Eickhoff et al., 2005) were used to pinpoint regions exhibiting significant interaction effects. Post-hoc group comparisons were conducted to further disentangle significant interaction effects. To this end, parameter estimates of atlas-based independent masks were extracted and subjected to post-hoc comparisons with FDR-correction in R-Studio (Benjamini and Hochberg, 1995). To further examine associations with current stress levels Spearman’s rank correlation were applied in SPSS to explore associations between extracted parameter estimates in significant cluster and PSS scores with a focus on participants without active treatment (ATDc-PLC).

## Results

### Mood state

Effects of treatment on mood were assessed using a mixed-design ANOVA model including mixture (ATD vs. ATDc) and intranasal spray (OXT vs. PLC) as between-subject factors, and timepoint (T1-T4; pre-oral administration, pre-intranasal administration, pre-fMRI, post-fMRI) as within-subject factor. In line with our previous study (Liu et al., 2021a), we observed significant main effects of timepoint on positive (*F*_(3,327)_ = 16.68, *p* < 0.001, η^2^_p_ = 0.13) and negative (*F*_(3,327)_ = 14.10, *p* < 0.001, η^2^_p_ = 0.12) affect, suggesting a general decline of positive and negative mood during the experiment (details see Supplementary **Table S2**). We also observed a trend for an interaction effect between of ATD and OXT on negative affect (*F*_(1,108)_ = 3.891, *p* = 0.051, η^2^_p_ = 0.035). Post-hoc comparisons revealed that participants who received ATD-OXT treatment reported a higher negative affect than who received ATD-PLC treatment (*p* = 0.043) or received ATDc-OXT treatment (*p* = 0.057, trend), while there was no significant difference between participants who received ATDc-PLC treatment and those received the ATD-PLC condition (*p* = 0.40) or the ATDc-OXT condition (*p* = 0.45), suggesting that the combination ATD and OXT treatment produced stronger increases on negative mood, while neither ATD nor OXT treatment did affect mood changes over the experiment in healthy individuals. No significant main effect of mixture or intranasal spray on both two types of mood was observed.

### Interaction between reduced serotonergic signaling and OXT on amygdala networks

In the whole-brain analysis, a significant interaction effect between treatments were observed with respect to left amygdala intrinsic coupling with a cluster located in the left hippocampus extending into the adjacent midbrain (MNI, *xyz* = [−20 −28 −10], *t*_*106*_ = 4.69, *p*_FWE-cluster_ = 0.046, *k* = 118 voxels, see **Figure 2A**). Probabilistic mapping by means of the Anatomy (version 2.2b) toolbox indicated that the interaction effect primarily encompassed the subiculum subregion of the left hippocampus. To disentangle the significant interaction effect, parameter estimates were extracted from the left subiculum using independently defined anatomical mask (Anatomy toolbox) for post-hoc comparisons which revealed that OXT (ATDc-OXT) significantly increased the connectivity relative to the PLC-reference (ATDc-PLC) group (*p*_FDR_ = 0.048), whereas pre-treatment with ATD (ATD-OXT) significantly attenuated the OXT-induced (ATDc-OXT) increase (*p*_FDR_ = 0.048) (see **Figure 2B**). Neither ATD (ATD-PLC) alone (*p*_FDR_ = 0.62) nor the combined application (ATD-OXT, *p*_FDR_ = 0.84) significantly increased amygdala connectivity in comparison to the PLC-treated reference (ATDc-PLC) group. To further determine which midbrain regions were included in the cluster we employed an atlas encompassing the corresponding regions (https://github.com/canlab/Neuroimaging_Pattern_Masks/tree/master/Atlases_and_parcellations/2018_Wager_combined_atlas) and observed that the cluster extended into the superior colliculus and dorsal part of the midbrain (see also **Figure S2**). No significant treatment interaction effects were observed for right amygdala intrinsic networks.

**Figure 2.**
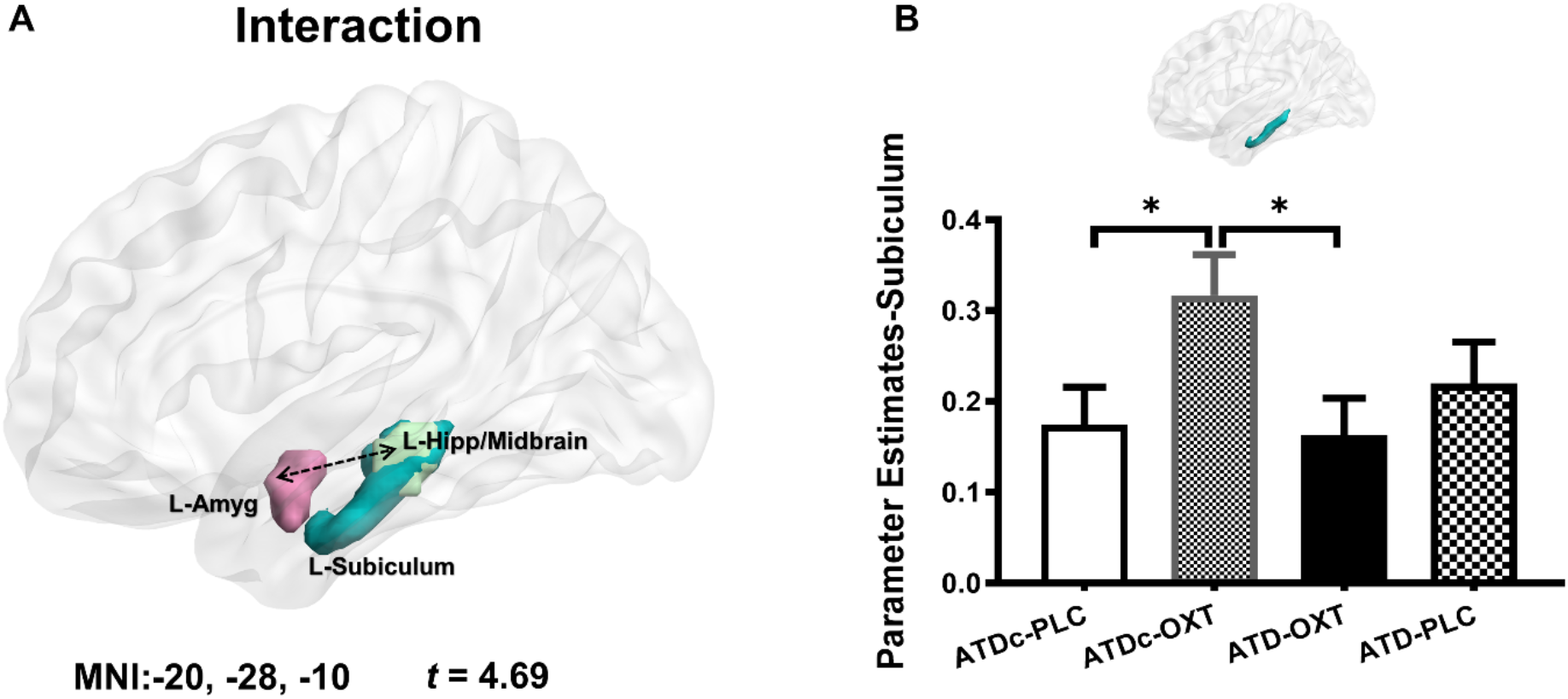
Interaction effects between oxytocin and serotonin on the intrinsic connectivity of the left amygdala. (A) The whole brain voxel-wise analysis revealed a significant negative interaction effect of treatments on the functional connectivity between the left amygdala and a left hippocampus/midbrain cluster (in light green, whole brain corrected at *p*_FWE-cluster_ < 0.05). The mask in light blue represents the left subiculum from the SPM Anatomy toolbox. (B) Post hoc comparisons between the treatment groups using extracted parameter estimates from the left subiculum anatomical mask. Group difference between ATDc-PLC and ATDc-OXT indicates that OXT significantly increased connectivity in this pathway in relative to the PLC-reference group, whereas the group difference between ATDc-OXT and ATD-OXT indicates that pre-treatment with acute tryptophan depletion significantly attenuated the OXT-induced increase. Bars indicate M ± SEM, * *p*_FDR_ < 0.05. Abbreviations: L, left; Amyg, amygdala; Hipp, hippocampus; OXT, Oxytocin; PLC, Placebo; ATD, Acute Tryptophan Depletion; ATDc, Acute Tryptophan Depletion control mixture.

### Associations with current stress

Associations between current stress and neural indices were examined by means of examining correlations between PSS scores and neural indices of significant clusters. Examining general associations of the identified pathway with current stress in the group without OXT or 5-HT modulation (ATDc-PLC) revealed a trend for a significant negative correlation between PSS scores and the left amygdala-left hippocampus/midbrain connectivity (*r*_*S*(26)_ = −0.34, p = 0.08, see **Figure 3**), suggesting that this pathway may be associated with current stress experience. Further control analyses in the groups receiving treatment did not reveal significant associations between this pathway and perceived stress (ATDc-OXT, *r*_*S*(24)_ = 0.25, *p* = 0.22; ATD-OXT, *r*_*S*(27)_= 0.10, *p* = 0.62; ATD-PLC, *r*_*S*(27)_ = −0.07, *p* = 0.72).

**Figure 3.**
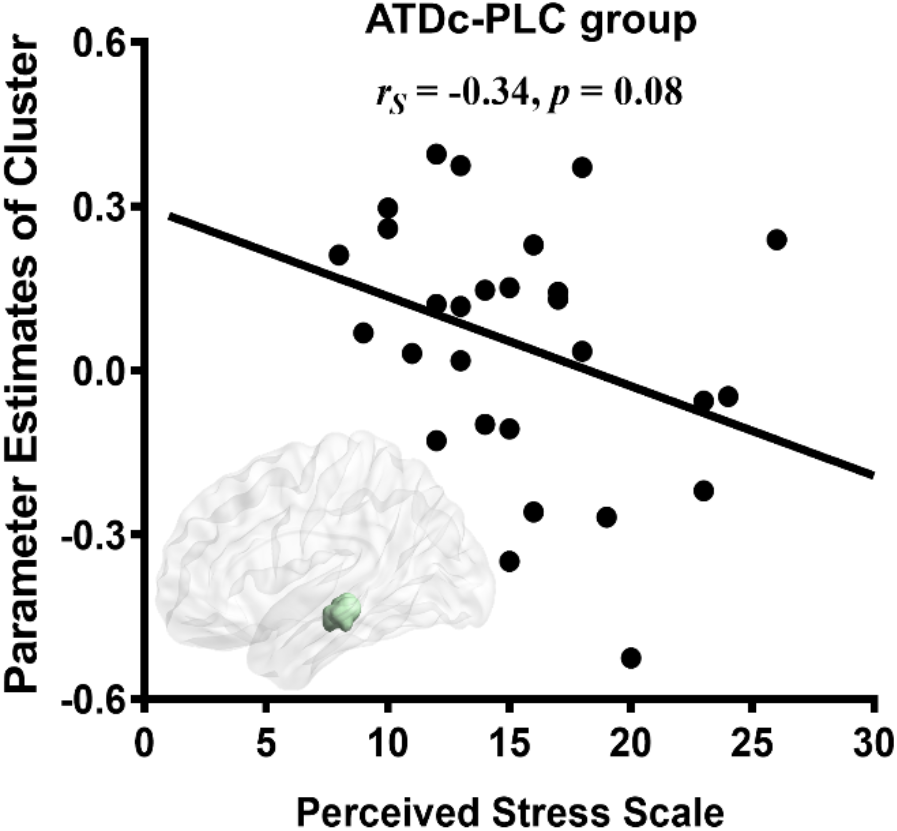
Associations of with currently experienced stress in ATDc-PLC group. Associations between Perceived Stress Scale scores and parameter estimates extracted from the amygdala-hippocampal pathway displaying correlation. Dots represent individual data. Abbreviations: PLC, Placebo; ATDc, Acute Tryptophan Depletion control mixture.

## Discussion

The present study aimed at determining interactive effects of the OXT and 5-HT system on intrinsic organization of the amygdala-centered networks. To this end, we employed a pharmaco-rsfMRI design that combined the administration of intranasal OXT with a transient decrease in central 5-HT signaling via acute tryptophan depletion or the respective control conditions and resting state fMRI. Examining treatments effects on the intrinsic amygdala networks revealed an interactive effect of OXT and 5-HT on the coupling of the left amygdala and a cluster encompassing the left subiculum of the hippocampus and extend to the left midbrain encompassing the superior colliculus. Post-hoc analyses convergently demonstrated that OXT increased intrinsic coupling in this pathway while this effect of OXT was significantly attenuated during transiently decreased central serotonergic signaling induced via ATD. In the placebo (ATDc-PLC) group lower connectivity in this pathway correlated with higher levels of experienced stress, suggesting a sensitivity of this pathway to current stress exposure. Together the present findings indicate that the effects of OXT on stress-associated amygdala intrinsic pathways with the hippocampus are critically mediated by 5-HT.

Previous pharmaco-rsfMRI studies have demonstrated that intranasal OXT modulates the connectivity of the amygdala with prefrontal regions (Dodhia et al., 2014; Koch et al., 2016; Eckstein et al., 2017; Jiang et al., 2021) and the hippocampus (Fan et al., 2015; Kirkby et al., 2018; Alaerts et al., 2019), while animal models and an initial human study suggest that some effects of OXT in the domains of anxiety and amygdala threat-reactivity are critically mediated by the 5-HT system (Yoshida et al., 2009; Mottolese et al., 2014). Within this context the present findings further extend previous findings that specifically the oxytocinergic modulation of the intrinsic coupling between the amygdala and the hippocampus/midbrain is critically mediated by interactive effects with the 5-HT system.

Across species, the amygdala-hippocampus-midbrain circuitry plays a role in the reactivity to potential threatening stimuli and stress reactivity (Belujon and Grace, 2011). In humans, intrinsic coupling in the amygdala-hippocampus pathway has been associated with acute and prolonged effects of stress exposure (Sripada et al., 2012; Vaisvaser et al., 2013; Fan et al., 2015; Zhang et al., 2016) and a recent intracranial electroencephalography study suggests that this pathway accurately tracks mood variations in humans (Kirkby et al., 2018). Whereas the functional role of the amygdala-midbrain pathways has not been extensively examined by means of rsfMRI in humans, animal models have documented its critical role in the selection of threat-induced behavioral responses and associated aversive learning processes (Steinberg et al., 2020). Previous intranasal OXT studies in humans reported an oxytocinergic modulation of amygdala-midbrain coupling during exposure to threatening social stimuli (Kirsch, 2005) as well as on oxytocinergic regulation of stress-induced changes in intrinsic amygdala-hippocampal coupling (Fan et al., 2015), suggesting a potential role of this pathway in the stress modulating effects of OXT. We found some evidence that the intrinsic communication in the amygdala-midbrain pathway is associated with current stress experience in the participants that did not receive active treatment, suggesting a role of the identified pathway in current stress experience. This may link the observed interactive effects to overarching theories on the regulatory role of OXT on stress- and anxiety-related processes (e.g. (Neumann and Slattery, 2016; Kendrick et al., 2018; Matsushita et al., 2019)).

The cluster exhibiting interactive effects of 5-HT and OXT encompassed the subiculum of hippocampus and the left superior colliculus and further dorsal parts of the midbrain. Previous rodent and nonhuman primates studies have suggested that both the subiculum and the superior colliculus are brain regions with particular dense OXT receptor expression (Elands et al., 1988; Shapiro and Insel, 1989; Freeman et al., 2014, 2014). The subiculum has been considered as stress-sensitive brain region in both animal and human studies (Mueller et al., 2004; Belujon and Grace, 2011; Teicher et al., 2012), and several studies reported an association between long term stress exposure and decreased the gray matter of this region (Teicher et al., 2012) and the amygdala (Lim et al., 2014). While previous studies reported strong functional connectivity between the amygdala and the hippocampus (Kiem et al., 2013), previous studies also reported that OXT can modulate this stress-related pathway (Fan et al., 2014, 2015; Hernández et al., 2015; Alaerts et al., 2019). The superior colliculus has been associated with defense reactions toward threat in rodent models (Coimbra et al., 2006), while such reflexive defense reactions were attenuated when the activation of amygdala and superior colliculus were inhibited simultaneously in non-human primates (Forcelli et al., 2016), suggesting a role of this pathway in defensive responses. In line with the functional role of this pathway previous OXT studies in humans reported that OXT induced a modulation effect on the functional connectivity between amygdala and the superior colliculus toward both threatening and non-threatening social stimuli (Kirsch, 2005; Gamer et al., 2010). Together, the current findings suggest that the effects of OXT on the intrinsic coupling between the amygdala with the subiculum and superior colliculus are critically modulated by interactive effects with the 5-HT system.

The interactive effects between the OXT and 5-HT system specifically affected the left amygdala intrinsic networks. Previous studies revealed inconsistent lateralization effects of OXT on the intrinsic amygdala networks in studies that examined the left and right amygdala separately (Sripada et al., 2013; Fan et al., 2014; Grace et al., 2019). In the previous literature such lateralization effects of OXT have been related to age, gender, stimuli type, or early life stress exposure (Striepens et al., 2012; Fan et al., 2015; Ebner et al., 2016; Grace et al., 2018, 2019; Alaerts et al., 2019). Left lateralized OXT effects have been repeatedly observed on cerebral blood flow activity and intrinsic connectivity of the left amygdala in young males (Paloyelis et al., 2016; Grace et al., 2019). Although the functional lateralization of the amygdala remains debated some early studies hypothesized that the left amygdala is stronger related to the conscious perception and regulation of emotional responses (Morris et al., 1998). Within this context the increased left amygdala-hippocampus/midbrain functional connectivity after OXT may reflect effects on threat perception and regulation (see also Liu et al., 2021).

We additionally observed an interaction effect between ATD and OXT on negative mood. In line with a number of previous studies ATD (Benkelfat et al., 1994; Evers et al., 2006; Talbot and Cooper, 2006) or OXT (Xu et al., 2019; Chen et al., 2020; Zhuang et al., 2021) per se did not affect mood. The interactive effect on negative mood was not predicted and may reflect complex interaction effects of 5-HT and OXT on rather non-specific negative affective states.

Findings of the present study need to be considered in the context of the following limitations. First, only male subjects were enrolled due to previous studies showing marked sex-differences in the 5-HT synthesis (Nishizawa et al., 1997) and sex-differences in the effects of OXT in rodents and humans (Dumais and Veenema, 2016; Dumais et al., 2017; Xu et al., 2020; Lieberz et al., 2020; Ma et al., 2018). Future studies need to determine whether the observed effects generalize to women. Second, tryptophan blood and OXT blood levels were not assessed. Although previous studies reported robust and selective decreases in 5-HT signaling (Crockett et al., 2008; Passamonti et al., 2012) and increases in (peripheral) OXT levels (Burri et al., 2008; Gossen et al., 2012) following similar the ATD or OXT treatment protocols as used in the present study, the additional examination of blood-level measures particularly in the combined treatment group may reveal important additional information on the interaction of the two systems and should be included in future studies. Finally, while the study involved a total of 112 participants in a complex pharmaco-fMRI design we did not include an a priori sample size calculation. The priori calculation for sample size and power in complex pharmaco-fMRI design is still limited and while our results are in line with our task-based findings in this sample demonstrating interactive OXT and 5-HT effects on threat-related amygdala processing (Liu et al., 2021a) the comparably low sample size in our study requires larger validation and replication studies (for details see also Cremers et al., 2017). While the present study focused on the connectivity of the amygdala as a single region, previous studies revealed subregional and nuclei specific effects of OXT on subcortical systems including the amygdala and basal ganglia (Eckstein et al., 2017; Zhao et al., 2019a; Martins et al., 2022) and future studies with larger samples may open the opportunity to examine OXT and 5-HT interaction effects on the subregional level. Overall, the present findings provide first evidence that effects of OXT on stress-associated amygdala-hippocampal-midbrain pathways are critically mediated by the 5-HT system in healthy men.

## Acknowledgments

We thank all subjects who participated in this study.

