## supplements for "Oxytocinergic modulation of stress-associated amygdala-hippocampus pathways in humans is mediated by serotonergic mechanisms"

**Authors:** Lan et al.

Benjamin Becker

The Clinical Hospital of Chengdu Brain Science Institute,

School of Life Science and Technology,

MOE Key Laboratory for Neuroinformation,

University of Electronic Science and Technology,

Xiyuan Avenue 2006, 611731 Chengdu, China

### Supplement materials

#### Supplementary Figures

**Figure S1** CONSORT flow diagram.

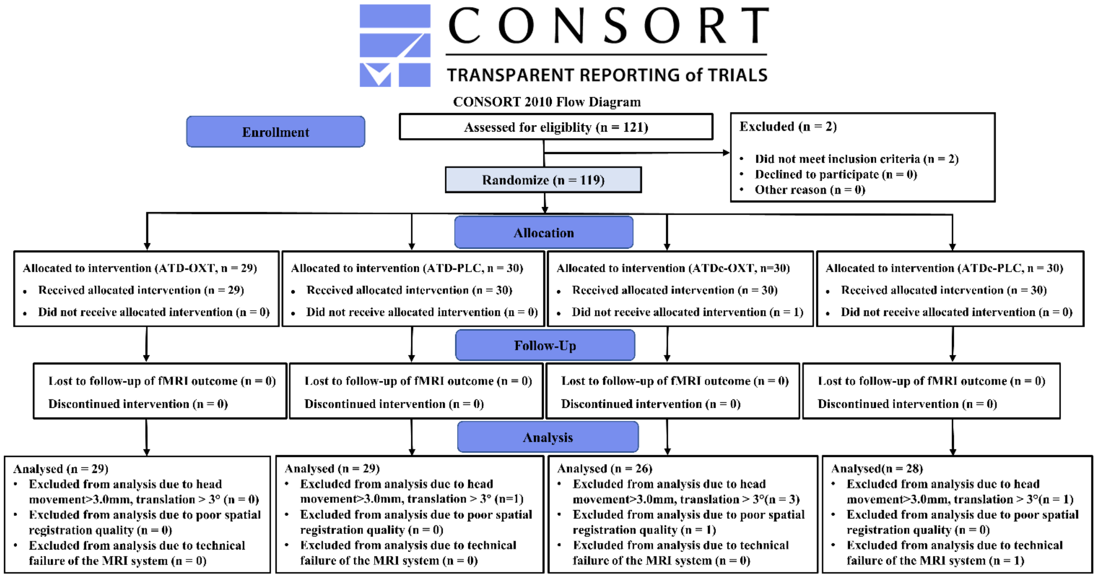


**Figure S2 Overlaps between observed cluster and superior colliculus.** The left hippocampus/midbrain cluster in yellow (whole brain corrected at $p_{\text{FWE-cluster}} < 0.05$). The mask in purple with 50% transparency represents the left superior colliculus from the Wager's brainstem atlas (https://github.com/canlab/Neuroimaging_Pattern_Masks/tree/master/Atlases_and_parcellations/2018_Wager_combined_atlas).

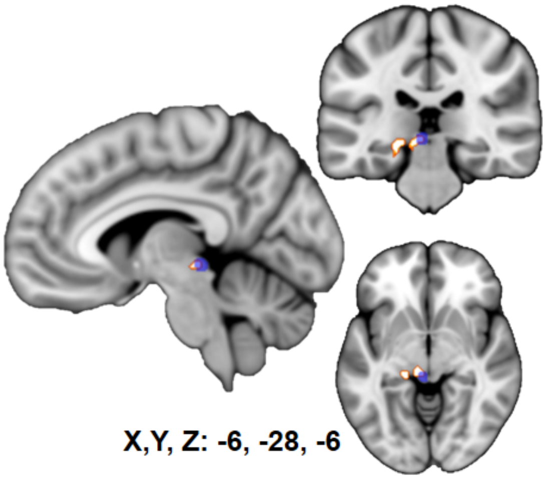

#### Supplementary tables

**Table S1** Components of the two amino acid mixtures in line with (Veen et al., 2007)

| Amino acids | ATD(g) | ATDc(g) |
|---|---|---|
| L-alanine | 4.1 | 4.1 |
| L-arginine | 3.7 | 3.7 |
| L-cysteine | 2 | 2 |
| L-glycine | 2.4 | 2.4 |
| L-histidine | 2.4 | 2.4 |
| L-isoleucine | 6 | 6 |
| L-leucine | 10.1 | 10.1 |
| L-lysine monohydrochloride | 6.7 | 6.7 |
| Methionine | 2.3 | 2.3 |
| L-phenylalanine | 4.3 | 4.3 |
| L-proline | 9.2 | 9.2 |
| L-serine | 5.2 | 5.2 |
| L-threonine | 4.9 | 4.9 |
| L-tyrosine | 5.2 | 5.2 |
| L-valine | 6.7 | 6.7 |
| L-tryptophan | 0 | 3 |
| Total | 75.2 | 78.2 |

**Table S2** Positive and negative mood assessment using PANAS in four groups

| Measurements | Time point | ATD-OXT n=29 | ATD-PLC n=29 | ATDc-OXT n=26 | ATDc-PLC n=28 |
|---|---|---|---|---|---|
| PANAS-P | Pre-oral | 24.9 (6.9) | 23.9 (8.3) | 24.3 (6.9) | 25.1 (8.2) |
| | Pre-intranasal | 22.1 (5.7) | 20.0 (6.7) | 21.4 (5.4) | 24.0 (6.7) |
| | Pre-fMRI | 21.3 (7.1) | 21.0 (7.9) | 20.2 (5.9) | 22.9 (6.6) |
| | Post-fMRI | 20.1 (6.1) | 19.8 (7.3) | 20.9 (6.0) | 23.2 (7.7) |
| PANAS-N | Pre-oral | 15.7 (9.0) | 13.1 (5.7) | 11.9 (5.2) | 13.3 (6.0) |
| | Pre-intranasal | 11.7 (3.7) | 11.2 (4.2) | 11.4 (5.4) | 12.2 (4.9) |
| | Pre-fMRI | 12.2 (4.3) | 9.6 (0.9) | 11.0 (4.9) | 10.9 (3.4) |
| | Post-fMRI | 12.9 (5.6) | 10.1 (1.5) | 10.0 (3.2) | 11.1 (4.5) |

Abbreviations: OXT, Oxytocin, PLC, Placebo, ATD, Acute Tryptophan Depletion; ATDc, Acute Tryptophan Depletion control mixture; PANAS-P, Positive and Negative Affect Schedule-Positive affect, PANAS-N, Positive and Negative Affect Schedule-Negative affect.
